# Gray matter cortical thickness predicts individual pain sensitivity: a multi-center machine learning approach

**DOI:** 10.1101/2022.06.14.496092

**Authors:** Raviteja Kotikalapudi, Balint Kincses, Matthias Zunhammer, Frederik Schlitt, Tobias Schmidt-Wilcke, Zsigmond T. Kincses, Livia Asan, Ulrike Bingel, Tamas Spisak

## Abstract

Pain sensitivity is known to considerably vary across individuals. While the variability in pain has been linked to structural neural correlates, it is still unclear how well these findings replicate in independent data and whether they are powerful enough to provide reliable pain sensitivity predictions on the individual level. In this study, we constructed a predictive model of pain sensitivity utilising structural MRI-based cortical thickness data from a multi-center dataset (3 centers, 131 healthy participants). Cross-validated estimates revealed a statistically significant and clinically relevant predictive performance (Pearson’s r = 0.36, p < 0.0005). The predictions were found to be specific to pain sensitivity and not biased towards potential confounding effects (e.g., anxiety, stress, depression, center-effects). Analysis of model coefficients suggests that the most robust cortical thickness predictors of pain sensitivity are the right rostral anterior cingulate gyrus, left parahippocampal gyrus and left temporal pole. Cortical thickness in these regions was negatively correlated to pain sensitivity. Our results can be considered as a proof-of-concept for the capacity of brain morphology to predict pain sensitivity, paving the way towards future multimodal brain-based biomarkers of pain.

**Highlights:**

1. We present a robust, brain structure-based multi-center predictive model for pain sensitivity.
2. Our model based on gray matter cortical thickness explains 13% of the variance in pain sensitivity and generalizes to out-of-center data.
3. The predictions are specific to pain sensitivity and not driven by potential confounders such as stress, depression, anxiety, and center-effects.
4. The most important predictors were rostral anterior cingulate cortex, parahippocampal gyrus and temporal pole, all negatively correlated with pain sensitivity.

## Introduction

Pain sensitivity is known to considerably vary across individuals (Nielsen et al., 2009). This variability has been reported to both forecast and accompany various clinical pain conditions (Meints et al., 2019; Nahman-Averbuch et al., 2019) and – besides peripheral aspects – it seems to be shaped to a large degree by individual differences of brain structure and function (Reddan and Wager, 2018; Spisak et al., 2020; Tu et al., 2019; Wager et al., 2013). Brain-based, objective markers of pain sensitivity are, therefore, of high interest for developing personalized diagnosis and treatment approaches (Tracey, 2021; Woo et al., 2017). Machine learning holds promise to exploit the rich information in magnetic resonance imaging (MRI) data to characterize pain robustly and objectively on the individual level. For instance, the neurologic pain signature (NPS) (Wager et al., 2013) utilizes task-based functional MRI data to objectively assess pain and has proven to be robust and generalizable to various pain conditions (Zunhammer et al., 2018). We have recently reported that resting state fMRI measurements also hold high predictive value in characterising pain; our Resting-state Pain sensitivity Network-signature (RPN) was found to robustly predict individual pain sensitivity on unseen data from different imaging centers (Spisak et al., 2020). These results clearly demonstrate the potential of functional MRI to serve as a basis for composite pain biomarkers (Tracey et al., 2019). Given the reliability of MRI-based brain morphological parameters (Iscan et al., 2015; Knussmann et al., 2022; Melzer et al., 2020) and the availability of anatomical imaging data in clinical routine, structural MRI-based pain sensitivity markers with sufficient predictive performance would hold promise for an even higher translational value than functional neuromarkers.

Although we still lack a rigorously validated structure-based predictive model for pain sensitivity, several studies have reported structural brain correlates of the sensitivity to pain on the group level. Voxel-based morphometry (VBM) studies have reported that pain sensitivity is significantly correlated to structures including the cingulate cortex (CC), precuneus, primary somatosensory cortex (S1), putamen, insula, parahippocampal gyrus (PHG), among others (Emerson et al., 2014; Neumann et al., 2021; Niddam et al., 2021; Ruscheweyh et al., 2018; Zhang et al., 2020). Surface based morphometry techniques (Fischl, 2012) can provide additional insights and, in contrast to VBM, a better account for gyrification via measures such as parcellated brain cortical thicknesses and volumes (Fischl et al., 2001). For instance, Erpelding et al., found a predominantly positive correlation between cortical thickness and heat pain thresholds in S1 (cold pain as well), posterior midcingulate cortex, and orbitofrontal cortex (Erpelding et al., 2012). Together, the studies have identified many possible neural correlates of pain sensitivity. However, it is still unclear whether these structural measures are robust enough to provide biomedically relevant predictions of pain sensitivity at the individual level. A recent study suggests that brain morphology (especially in the prefrontal cortex) may have the potential for predicting an individual’s pain sensitivity (Zou et al., 2021), as demonstrated for pain sensitivity to laser, heat and cold pain using T_1_-based indices of morphological connectivity (how similar the morphology of two regions are; (Tijms et al., 2012)). However, Zou et al. did not systematically evaluate the specificity of their model to pain sensitivity (by evaluating potential confounding effects (Dinga et al., 2020; Giusti et al., 2021; Klauenberg et al., 2008; Sivertsen et al., 2015). Therefore, although their results are generally promising regarding the potential of brain structure to predict individual pain sensitivity, the specificity and the neurobiological validity of their predictive model cannot be thoroughly evaluated. Moreover, due to the single-center design and the lack of unbiased estimates of predictive performance in this study (i.e., the lack of nested-cross validation), it remains unclear, whether brain morphology can provide sufficient information for clinically relevant individual-level pain sensitivity predictions.

Here we aim to develop a brain morphology-based predictive model that is (i) specific to individual pain sensitivity and not confounded by other related but out-of-interest factors, (ii) generalizable for unseen data, regardless of imaging details (site, scanner, sequence parameters) and (iii) explainable, i.e., it provides insights into the underlying mechanisms. To be able to evaluate the specificity and neurobiological validity of our model (i), we assess a comprehensive set of potential confounder variables and analyze their effect on the model predictions with a dedicated statistical procedure (Spisak, 2021). To avoid the ‘standardization fallacy’ (Voelkl et al., 2021) and to ensure that the model is robust for differences in image acquisition details and center effects (ii), we train the model on multi-center data and use a balanced nested-cross validation scheme and a clean, minimalist workflow (with a low number of free parameters) to avoid “vibration effects” (Varoquaux, 2018) and “feature leakage” (Mateos-Pérez et al., 2018) during the machine learning procedure and to obtain unbiased estimates of predictive performance. To ensure model explainability (iii), we keep the model structure as simple as possible and restrict the analysis to regional cortical thickness estimates which allow for a quantitative characterization of effect sizes.

## Methods

### Participants

A combined total of n = 133 participants were recruited, and experimental procedures were carried out at three different study sites, namely Ruhr University Bochum – Germany (study 1, n=39), University Hospital Essen – Germany (study 2, n=49), and University of Szeged – Hungary (study 3, n=45). Reimbursement was done only for study 1 and study 2 at 20 €/h. All studies were conducted in accordance with the Declaration of Helsinki, and ethically approved by the local or national committees (Register Numbers: 4974-14, 18-8020-BO, 057617/2015/OTIG, and ETT TUKEB for studies 1-3, respectively). For inclusion-exclusion criteria in detail, please refer to *table 3* from Spisak et al. (Spisak et al., 2020). In brief, inclusion criteria comprised of no chronic disease condition, age group of 18 to 40 years (target age = 25y), right-handed non-smoking participants, and balanced sex distribution. Moreover, participants were excluded in presence of acute or chronic neurological endocrine, or psychiatric conditions, acute infections, use of psychotropic or analgesic based substances, wounds, scars, or skin conditions which could potentially affect the pain stimuli-based experimental setup. MRI scans underwent a quality check procedure to exclude images with incomplete whole brain coverage or motion artefacts. Overall, 2 MRI scan with an incomplete frontal brain acquisition was excluded from final analysis (n = 131).

### Pain sensitivity measure

Heat (HPT), cold (CPT) and mechanical pain thresholds (MPT) were acquired from the participants following the established quantitative sensory protocol (QST) (Rolke et al., 2006). While warmth (WDT) and cold detection thresholds (CDT) were collected in study 1 and study 2, in addition, mechanical (MDT) thresholds were also collected for study 3. Thermal thresholds were obtained using ATS thermodes, on a skin surface 30 × 30 mm, at the left palmar forearm, proximal to the wrist crest. For thermal stimulators, MSA thermal stimulator (Somedic, Horby, Sweden) was used in study 1, and Pathway thermal stimulator (Medoc Ltd., Ramat Yishai, Israel) was used in study 2 and 3. Increasing and decreasing thermal thresholds were applied to the skin, and the baseline temperature was kept at 32 degrees Celsius. Using a button press, participants indicated heat and cold pain onsets. Instead of 3 (original protocol), 6 stimuli repetitions were conducted for all thermal thresholds to address between-subject variance. For calculating HPT and CPT, arithmetic mean of measurements (2 to 6) was taken, excluding the first measurement (test measure) for each participant. For mechanical stimuli, alternatively 5 increasing (for MPT) and 5 decreasing (for MDT) trains of pinprick (MRC Systems, Heidelberg, Germany) stimuli were applied to the participants. Participants were given instructions to categorize the stimuli as either noxious, or non-noxious. Spisak et al., also provides the details regarding the experimental paradigm (Spisak et al., 2020). MPT and MDT measures were calculated as the log-transformed geometric means of all mechanical pain and detection measures. Finally, QST was calculated as a composite score composed of HPT, CPT, and MPT. All three measures were z-transformed within each center, and HPT, and MPT were inverted (× -1; the algebric signs ‘-’ were adjusted so that it reflects the overall pain sensitivity of the participant in the final composite score, i.e., QST). This was followed by calculating the arithmetic mean of the three pain thresholds for each participant. QST measures that were at least 2.5 standard deviations from mean were excluded. None of the QST measures were identified as outliers.

### Additional measures

We acquired additional measures wherever possible for the three study centers to evaluate potential confounders. Additional measures included age, body mass index (BMI), date of the first day of last menses (females), level of education (primary, secondary, and university), self-rated pain sensitivity questionnaire: PSQ (Ruscheweyh et al., 2009), pain catastrophizing scale: PCS (Sullivan et al., 1995), the Pittsburgh Sleep Quality Index: PSQI (Buysse et al., 1989), perceived stress questionnaire: PSQ20 (Levenstein et al., 1993), state-trait anxiety inventory: STAI (Spielberger, 1983), short German version of the depression scale: ADS-K (Lehr et al., 2008). Blood pressure was recorded both prior to the MRI and the pain sensitivity experiment. In study 1, the so-called T_50_ values i.e., the temperature value where participants showed a heat pain rating of 50 on a visual analogue scale from 0 – 100 (0 = no pain, 100 intolerable pain), were also measured. Please refer to **supplementary table 1** for study specific sample sizes.

### MRI data

Our study utilizes 3D T1-weighted (structural) images in all study centers, performed on 3T scanners with an isotropic voxel size of 1 mm^3^. To avoid the “standardization fallacy” (Voelkl et al. 2021), no further harmonization of imaging sequences has been performed across sites. Site specific scanning parameters are listed in in **table 1**. The three centers utilized different scanners, namely Philips Achieva (study 1), Siemens Magnetom Skyra (study 2) and GE Discovery 750 w MR (study 3). Structural image processing of T_1_-weighted images was facilitated by freesurfer software version 6.0 available at https://surfer.nmr.mgh.harvard.edu/. The underlying working framework for image processing using freesurfer is covered extensively elsewhere (Collins et al., 1994; Dale et al., 1999; Fischl and Dale, 2000; Fischl et al., 2001; Fischl et al., 2002; Fischl et al., 1999a; Fischl et al., 1999b; Fischl et al., 2004). To this end, we decided to include gray matter thickness alone compared to other measures of morphology (like GM volume, area and density or morphological connectivity). The choice of gray matter thickness was motivated by the reliability and the quantitative nature of this measure (Han et al., 2006), as well as its potential independence from factors such as brain surface area and head size in comparison to GM volume, where factor correction procedures are not well standardized and could induce unnecessary statistical artifacts (Schwarz et al., 2016). Moreover, cortical thickness as a predictor of pain sensitivity holds promise for a high neurobiological relevance as it has already been reported to be associated with pain sensitivity by several studies (Erpelding et al., 2012; Grant et al., 2010). The steps of processes involved in obtaining cortical thickness using *recon-all* measures are listed with a brief explanation for each process at https://surfer.nmr.mgh.harvard.edu/fswiki/recon-all. Briefly, native space images were transformed to a standard space with a Talairach transformation (Talairach, 1988). Next image processing steps included correction for motion artifacts (Reuter et al., 2010), skull stripping (Ségonne et al., 2004), removal of cerebellum and brain stem, correction of non-uniform intensities (Sled et al., 1998), segmentation of subcortical and white matter (WM) and deep gray matter (GM) regions that helps in estimating GM-WM junction, tessellation of GM-WM boundary, smoothing of tessellated surface, automated topology correction (Ségonne et al., 2007), and generating a model for pial surface (GM-cerebrospinal fluid junction, (Dale et al., 1999; Fischl and Dale, 2000)). On the basis of Desikan-Killiany atlas (Desikan et al., 2006), cortical thickness was measured as the average of distance to the nearest point on the pial surface from each vertex of the tessellated WM boundary, and from that point back to the nearest point on the WM boundary. This procedure for obtaining cortical thicknesses has been validated against histological and manual measurements, and its reliability has been established across different scanning parameters such as MRI sequences and scanning machines (Jovicich et al., 2009; Rosas et al., 2002; Salat et al., 2004). We considered the cortical thickness measures from the freesurfer analysis, which resulted in a total of 68 regional thickness measures (34 per hemisphere, measured in millilitres).

**Table 1.** MRI acquisition information

|  | Study 1 | Study 2 | Study 3 |
| --- | --- | --- | --- |
| **Scanner** | Philips Achieva X | Siemens Magnetom Skyra | GE Discovery MR750w |
| **Head coil** | 32-channel | 32-channel | 20-channel |
| **Sequence** | MPRAGE | MPRAGE | IR-FSPGR |
| **FOV** | 256 × 256 × 220 mm<sup>3</sup> | 256 × 256 × 192 mm<sup>3</sup> | 256 × 256 × 172 mm<sup>3</sup> |
| **TR** | 8500 ms | 2300 ms | 5.3 ms |
| **TE** | 3.9 ms | 2.07 ms | 2.1 ms |

Study specific MRI acquisition information is presented including scanner protocols.

### Feature harmonization

The feature set comprised 68 features (cortical thickness in 68 regions of the Desikan-Killiany atlas (Desikan et al., 2006)) for 131 subjects. Regional cortical thickness measures were scaled by normalizing with mean cortical thickness at the participant-level. As cortical thickness measures were previously reported to be highly specific to scanning sites (Fortin et al. 2018), the feature set was harmonized to address the site-effects. The harmonization process was performed with ComBat, a batch-effect correction tool frequently used in genomics (Fortin et al., 2018). ComBat was configured to mitigate site effects, while still preserving the biologically relevant effects of age and sex. To avoid feature leakage (Mateos-Pérez et al., 2018), harmonization (with preserving for age and sex) has been incorporated into the machine learning cross-validation framework, detailed in the next paragraph (Machine learning pipeline). Leakage was prevented by using two functions: ComBat *fit* and ComBat *transform*. The fit function can be used on the (outer) training set to fit a ComBat model, whereas the transform function can be used to simply apply the already fitted ComBat transformation on the hold-out feature set. As a result, the transformed train set is guaranteed not to learn any information from its corresponding test set, thus avoiding data leakage which could potentially bias the model predictions.

### Machine learning pipeline

The machine learning (ML) framework is depicted in **figure 1**. First, the entire sample (feature set: 132 × 68 and corresponding target set = QST scores, 132 × 1) was split into *train* (feature set + target) and its corresponding *test* (only feature set) sets using a balanced 10-fold cross validation (CV, using GroupKFold: 1-fold corresponds to 1 iteration), where each fold would hold approximately the same amount of data from all three samples. This CV framework would serve as the outer loop for our ML pipeline. In this outer loop, the train data is fit + transformed using ComBat, and the test set is transformed with this already-fitted ComBat model, as described above. In the inner CV-loop, the training set from the outer loop is further split into a train subset and a validation subset in a 10-fold CV fashion. Subsequently, in each fold, the transformed train subset with its corresponding target values (i.e. QST-based composite pain sensitivity scores) was used to fit a linear regression model with least absolute shrinkage and selection operator: LASSO (Tibshirani, 1996) via the scikit-learn (Pedregosa et al., 2011) package (https://scikit-learn.org/stable/). LASSO shrinks the features of less importance and, potentially, eliminating them (nullifying coefficient/weight of the feature), to reduce model complexity and prevent overfitting. LASSO’s tendency to fully eliminate unimportant features can also improve the interpretability of the resulting model. Shrinkage in LASSO is implemented by an L1-penalty equal to the absolute value of the magnitude of feature coefficient and can be adjusted through tuning a single hyperparameter: α (learning rate). At different hyperparameter values for α ([0.0001, 0.001, 0.01, 0.1, 1, 10, 1000, 10000]), the LASSO model was fit to the inner train set and evaluated on the validation subset in terms of mean squared error (MSE) when predicting the composite pain sensitivity scores. Best LASSO model estimator in each inner loop was applied to create predictions for the actual outer test set (final predictions), in each of the 10 outer CV loops. Total intracranial volume was regressed out from the final predictions (Dinga et al., 2020), using a linear model that was fit on the train predictions, and later applied on the predictions of the test sets in each of the 10 CV iterations. We evaluated feature importance for each “best estimator model”, corresponding to the 10 outer cv iterations and considered features which have a non-zero coefficient, to be robust predictors for pain sensitivity. Pearson’s *r* was calculated for all associations and its p-value was calculated using non-parametric permutation testing with 10, 000 permutations.

**Figure 1.**
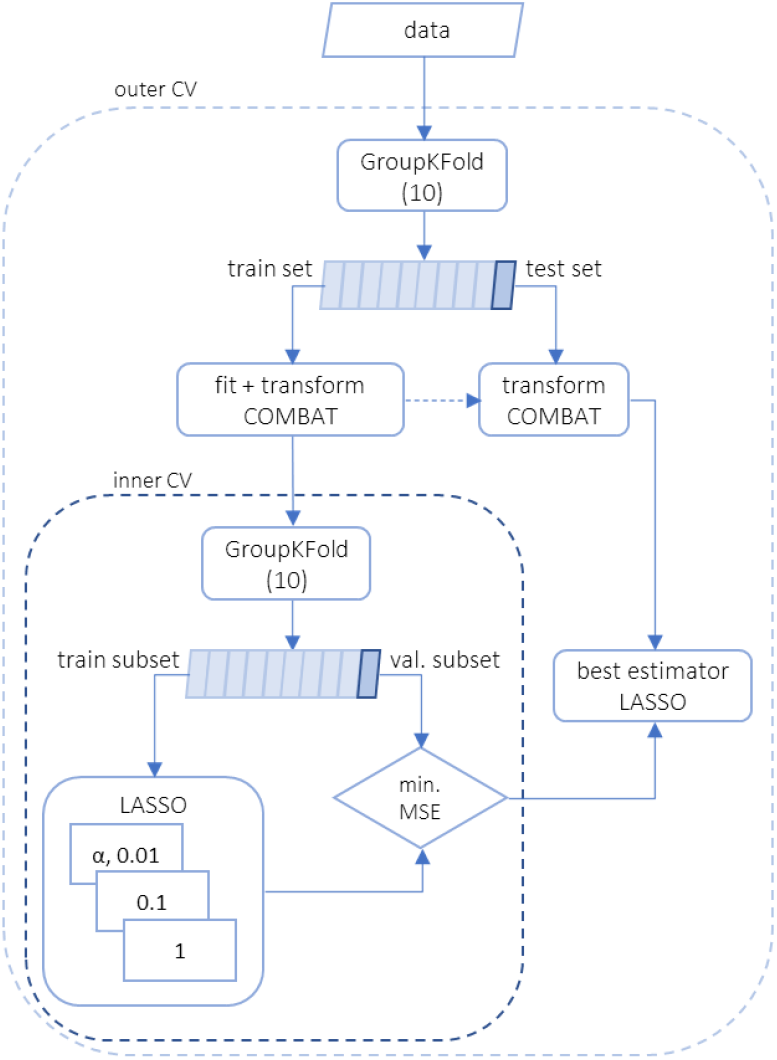
Flow chart of the machine learning pipeline. The flowchart shows splitting multi-center data into train and test sets and subsequent processes. In the outer cross-validation (cv), train sets consist of participant’s cortical thickness matrix and QST scores, which are used for fitting LASSO models. In the test sets the participants’ cortical thickness measures are fed into the already fitted LASSO model (optimized in the inner cv loop) to provide cross-validated predictions for the pain sensitivity scores. The nested cross-validation scheme and the use of the fit-transform scheme for data harmonization (ComBat) avoids feature leakage and provides unbiased estimates of predictive performance. The weight signs (+/-) would determine the directionality of association (positive or negative) of regional cortical thickness with pain sensitivity. Whereas the absolute magnitude of weight would determine the rank (effect size) of the region in predicting pain sensitivity (highest magnitude will have a rank = 1).

### Confounder effects

To evaluate potential confounding bias in the model predictions for center, demographic and psychological based additional measures, we used mlconfound as implemented in the python-package available from https://mlconfound.readthedocs.io (Spisak, 2021). We used the partial confounder test with the null hypothesis of ‘no confounder bias’, which is tested by probing the conditional independence of the predicted pain sensitivity on each additional measure, given the observed pain sensitivity. The test is distinguished from alternative approaches by its robustness to non-normally and non-linearly dependent predictions, therefore it is applicable without investigating the conditional distribution of the predictions on the target and the confounder for normality and linearity. A p-value < 0.05 implies confounding bias (i.e., the predictions are driven partially by the confounder).

### Leave-one-study-out

Along with the main analysis, we also assessed out-of-center generalizability by means of a leave-one-study-out cross-validation, that is, predicting each study, by fitting the above-described machine learning model on the remaining two studies. Pain sensitivity in the left-out center was predicted with the model that performed best on data from the remaining two centers (with hyperparameter α optimized in a nested cross-validation loop). That is, study 1+2 (1+3 and 2+3) was used to estimate pain sensitivity in a completely unseen study 3 (2 and 1). Parameters such as α-values, innerCV data splitting scheme (GroupKFold(10), remained the same as mentioned in the previous section (machine learning pipeline). It should be noted that, since ComBat can only consider batch IDs that it has already seen during training, the final predictions were performed twice, i.e., the left-out study was considered by combat to be from an identical source as either the first or the second from the training studies (and mean predictions were considered). All analyses (multi-center cortical thickness data and codes) are available as a repository on GitHub at https://github.com/rkotikalapudi/gmCT-predictive-modelling.

### HCP1200 analysis

To further characterise the model, as well as its most important predictors, in terms of specificity to pain sensitivity, we analysed already processed freesurfer data from the Human Connectome Project (www.humanconnectome.org, release data March 01, 2017). While the HCP1200 project does not involve measures of pain sensitivity, it contains two pain related scores, namely, the pain intensity and pain interference surveys (both self-reported). As these measures are only indirectly related to the composite pain sensitivity measure of our study, any significant association can be considered as a sign of the out-of-context generalizability of our model (or its predictors). Conversely, due to the large sample size (N=1095) and high statistical power, the lack of significant association strongly suggests that the model (or the predictors) is specific to QST-based pain sensitivity and do not generalize to self-reported pain phenotypes of the HCP data. Associations were tested with Pearson’s correlation. We tested only those regions that emerged as the non-zero weighted features (a priori) in our LASSO driven predictive model. The correlations’ one-sided p-value was obtained using non-parametric permutation testing (n = 10,000). Like our data analysis, the cortical thickness values were normalized with mean thickness and area measures with eTIV. Prior to performing the correlations, cortical measures were adjusted for age, sex, TIV and batch effects using ComBat.

## Results

### Model predictions

The multi-center model predicted pain sensitivity in unseen participants with a medium effect size, Person’s correlation r = 0.36 (R^2^ = 0.13) at p-value = 0.0002. The predicted values also correlated significantly with QST scores from each individual site, i.e. study 1 (r, p = 0.42, 0.01, **see supplementary figure 1 a-c**), study 2 (0.32, 0.01) and study 3 (0.35, 0.01). Multi-center model predictions for pain sensitivity are presented on **figure 2a**.

**Figure 2.**
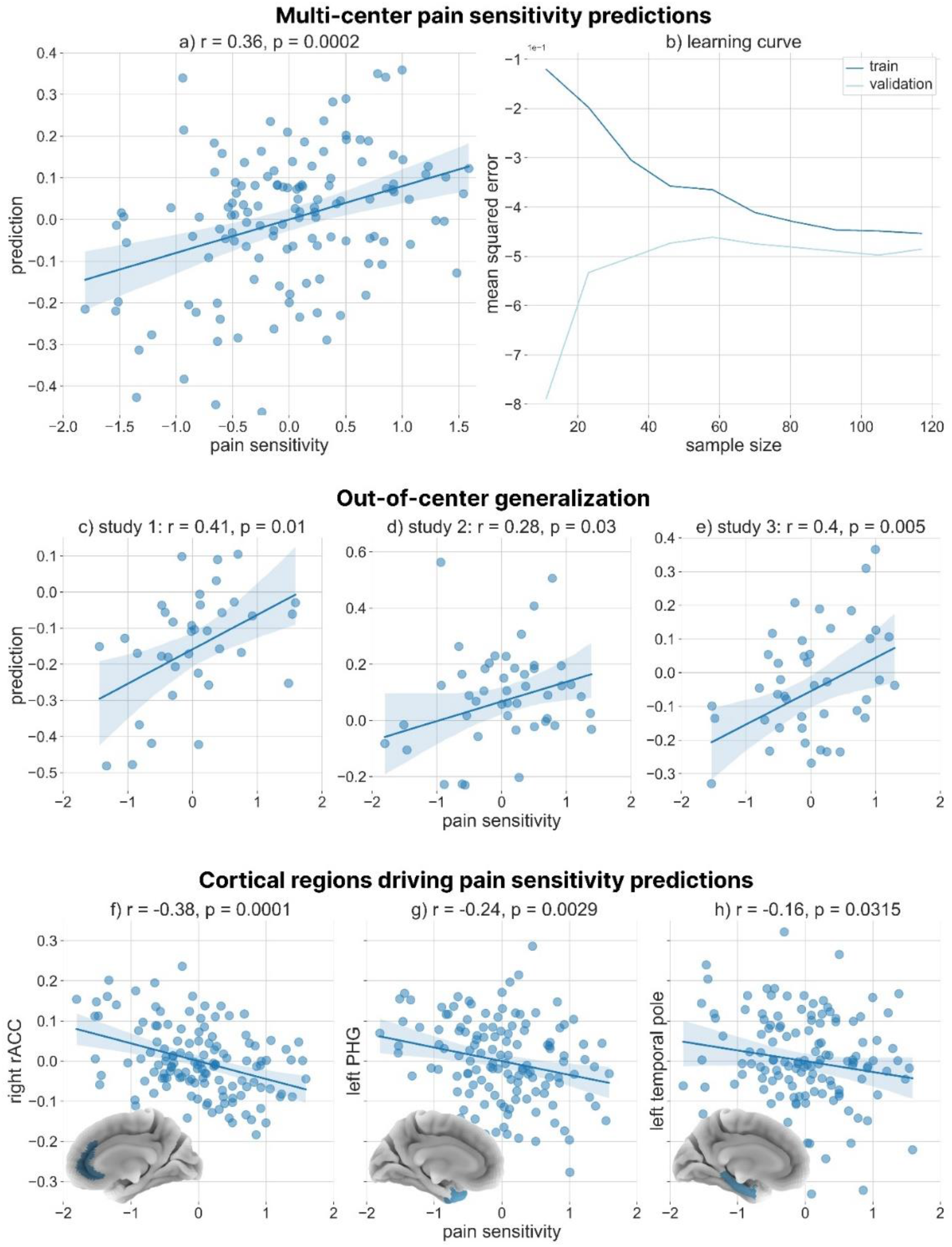
Predictive model outcomes. Scatter plots and regression lines (with 95% confidence intervals) depicting the correlation between pain sensitivity and the machine learning model’s predictions are shown for a) the pooled multi-center dataset (balanced 10-fold cross-validation) and for leave-one-sample-out analysis: c) study 1, d) study 2 and e) study 3. b) Learning curve for the best model for the train and validation sets. The learning of the model saturates approximately at sample size > 60. Scatter plots and regression lines (with 95% confidence intervals) depicting the correlations between individual pain sensitivity and cortical thickness of the pain predictive regions i.e., f) rACC, g) PHG and h) TP are shown for the pooled multi-center data.

### Model properties

The best model was found with an α = 0.01 (hyperparameter), consequently across all 10 iterations of the outer loop cross-validation in the multi-center analysis. The applied machine learning model (LASSO) eliminates features of less importance by assigning a feature weight (coefficient) = 0. Hence, we interpret the features with non-zero weights (coefficients) as the most prominent regions involved in predicting pain sensitivity. There were 6 regions that had non-zero coefficients in at least one of the cross-validation iterations and 3 out of them emerged in all iterations (**table 2**). These three regions also displayed the highest predictive importance (mean predictive coefficient, averaged over cross-validation iterations) in the model. Based on absolute weights (coefficients of the feature), the right rostral anterior cingulate cortex (right rACC) provided the highest contribution to the model’s predictions, which was followed by left parahippocampal gyrus (left PHG) and left temporal pole (left TP). All three features were negatively associated with the pain sensitivity scores (**figure 2 f-h**) i.e., right rACC (r = −0.38, p = 0.0001), left PHG (−0.24, 0.003) and TP (−0.16, 0.03). Three more regions, the right PHG (−0.22, 0.01), right frontal pole (0.12, 0.09), and left entorhinal cortex (−0.13, 0.07), have also emerged with non-zero weights it at least one CV iteration, but their appearance was not consistent (they emerged only in 1, 5 and 2 CV iterations, respectively).

**Table 2.** Cross-validation feature information

|  | right rACC | left PHG | left TP | right FP | right PHG | left ENT |
| --- | --- | --- | --- | --- | --- | --- |
| mean weight | -1.67 | -0.53 | -0.21 | 0.11 | -0.02 | -0.01 |
| STD | 0.24 | 0.18 | 0.10 | 0.11 | - | 0.001 |
| frequency (/total CV) | 10/10 | 10/10 | 10/10 | 5/10 | 1/10 | 2/10 |
| feature rank | 1 | 2 | 3 | - | - | - |

Table depicts the robustness of predictive features across cross validation folds. Mean and standard deviation of LASSO model coefficients across all outer cv iterations are shown for all features that were given a non-zero coefficient at least in one cross validation fold. The frequency of occurrence in the model across all cv-iterations reveals that the right rACC, the left PHG and the left TP are the most robust predictors of pain sensitivity. The hyper-parameter $\alpha$ remained constant across all 10 cross-validations at 0.01. ENT stands for entorhinal cortex.

### Confounder analysis

The cofounder analysis revealed no significant (p-value < 0.05) confounding bias for any of the investigated demographic, physiological and psychological variables (**table 3**).

**Table 3.** Confounder analysis for prediction results with additional measures

| No. | Additional measures | Correlation with QST (r) | Correlation with pred. QST (r) | Significance of confounder bias (p) |
| --- | --- | --- | --- | --- |
| 1 | BP MRI systole | -0.11 | -0.07 | 0.53 |
| 2 | BP MRI diastole | 0.01 | 0.13 | 0.20 |
| 3 | BP QST systole | -0.1 | -0.03 | 0.82 |
| 4 | BP QST diastole | -0.05 | 0.04 | 0.72 |
| 5 | MRI QST diff. | 0.06 | 0.07 | 0.48 |
| 6 | sex | -0.09 | -0.02 | 0.88 |
| 7 | day of menses | -0.09 | 0.05 | 0.72 |
| 8 | age | 0.03 | -0.07 | 0.42 |
| 9 | BMI | 0.01 | 0.04 | 0.71 |
| 10 | education | -0.05 | -0.13 | 0.26 |
| 11 | alcohol per unit | 0.15 | 0.21 | 0.25 |
| 12 | alcohol per week | -0.05 | -0.05 | 0.62 |
| 13 | PCS: catastrophizing | 0.09 | 0.03 | 0.72 |
| 14 | PCS: rumination | 0.14 | 0.01 | 0.93 |
| 15 | T50 | -0.45 | 0.12 | 0.71 |
| 16 | GLX | 0.36 | 0.37 | 0.10 |
| 17 | GABA | 0.24 | 0.11 | 0.58 |
| 18 | anxiety state | -0.01 | 0.03 | 0.75 |
| 19 | anxiety trait | 0.11 | -0.02 | 0.81 |
| 20 | pain questionnaire | 0.16 | 0.03 | 0.74 |
| 21 | depression | -0.11 | -0.01 | 0.91 |
| 22 | stress | 0.17 | 0.06 | 0.64 |
| 23 | sleep quality | -0.04 | 0.1 | 0.42 |
| 24 | CDT | 0 | 0.07 | 0.50 |
| 25 | WDT | 0 | 0.08 | 0.44 |
| 26 | MDT | 0.05 | 0.02 | 0.32 |
| 27 | study center | -0.01 | 0.04 | 0.65 |
| 28 | Estimated TIV | -0.02 | -0.02 | 0.842 |

Confounder analysis was performed between the predicted pain sensitivity, each of the potential confounder (1 additional measure at a time), given the observed pain sensitivity using mlconfound toolbox (Spisak 2021). A p-value < 0.05 suggests that confounder bias exists, and the predictions are not exclusively driven by the cortical thickness, but also by the additional measure.

### Out-of-sample generalization

The results of the leave-one-study-out analysis are presented on **figure 2 (c-e)** and **supplementary figure 1**, with coefficients of predictive regions in **table 4**. In brief, the predictions across all three studies were significant at p < 0.05. Common predictor across all the studies was the right rACC. Left PHG was an additional common predictor for study 1 and 3 (right PHG in case of study 2), left TP for study 2 and 3. Other predictors included right FP (study 1) and left entorhinal cortex (study 3).

**Table 4.** Leave-one-study-out analysis results.

| Left-out-study-ID | weights |
| --- | --- |
| Study 1 | right rACC = - 1.72, left PHG = - 0.49, right FP = 0.14 |
| Study 2 | right rACC = - 1.39, left TP = - 0.89, right PHG = - 0.87 |
| Study 3 | right rACC = - 1.86, left PHG = - 0.77, left ENT = - 0.29, left TP = - 0.06 |

Regularization factor $\alpha$ remained constant in all validations at 0.01. Study 1 (2 and 3) was a completely new data to the model that was calculated using study 2 - 3 (1 - 3, 1 - 2). TIV was regressed out from final predictions within the cross-validation framework. Age, gender was adjusted along with batch (study center) in the ComBat model prior to predictions. ENT stands for entorhinal cortex.

### Generalizability to self-reported pain measures in the HCP1200 dataset

The proposed model was found to be specific to QST-based pain sensitivity as its output was not found to be significantly associated with self-reported pain intensity and inference in the HCP dataset. We also checked for associations between morphological measures (cortical thickness and area) of the three most predictive regions (rACC, TP, PHC) and pain-related questionnaires (pain intensity and interference ratings) in the HCP1200 dataset (**supplementary table 2**). There were no associations found for pain with rACC. There was a weak but significant negative correlation between the thickness of TP and pain intensity score (r = −0.06, p = 0.02), observed on both sides. For pain interference T-score, there was also a weak but significant negative correlation with the area of PHG (r = −0.06, p = 0.02), bilaterally.

## Discussion

In this study, we have developed a multi-center predictive model for pain sensitivity based on structural brain morphology. Our study has two main findings. First, the proposed model can be considered as a proof-of-concept for the capacity of brain morphology to yield robust and specific individual-level predictions for pain sensitivity, posing brain structure as a promising component of future multimodal brain-based biomarkers of pain. Second, our results broaden our knowledge about the brain structural correlates of the individual differences and alterations in pain sensitivity by identifying three key regions that drive the morphology-based pain sensitivity predictions, the right rostral anterior cingulate cortex (rACC), the left parahippocampal gyrus (PHG), and the left temporal pole (TP).

When evaluating the potential of the proposed model for brain-based quantification of pain sensitivity, one of the most important considerations is the expected accuracy of the model’s predictions. In our study, grey matter cortical thickness explained 13% of the variance in pain sensitivity (r=0.36 for single model based on all centers and r=0.28-0.41 in the leave-one-center-out cross-validation). The achieved medium-range predictive effect size (according to Cohen’s recommendations) may hold clinical relevance (Dworkin et al., 2008) and renders brain morphology as promising modality for the objective brain-based characterisation of pain.

We have multiple reasons to believe that the reported performance estimates are realistic and will generalize well to independent data in future external validation studies. First, all predictions were computed inside a leakage-free nested cross-validation framework with a minimal, conventional feature preprocessing approach to avoid effect size inflation due to methodological choices, a.k.a “vibration effects” (Varoquaux, 2018). Second, in both the multi-center cross-validation and the leave-one-study-out analysis, the parameters of the predictive model exhibited a considerable robustness. Namely, the hyperparameter α remained constant in all cases and the key predictors i.e., rACC, PHG and TP emerged in the same order-of-importance in all iterations in the multi-center training. Third, given the heterogeneity between the included datasets (different scanners, sequences, research staff, etc.), the leave-one-study-out analysis strongly suggests that the proposed model can be expected to generalize well to data from new study centers.

Biomedically relevant predictive performance and generalizability to new data are necessary but not sufficient criteria for brain-based biomarker candidates (Spisak, 2021; Woo et al., 2017). Specificity to pain and the absence of bias towards potential confounders are also crucial factors that determine the clinical and translational of utility of brain-based predictive models of pain (Spisak, 2021). Our confounder analysis showed that the predictions of the proposed model are not confounded by variables-of-no-interest - such as center effects (which have been shown be detrimental for cortical thickness calculations (Fortin et al., 2018)), demographics, total intracranial volume, blood pressure, menstrual cycle, alcohol consumption or sleep quality. The predictions were also not primarily driven by sensory detection thresholds (as measured with QST) suggesting that the model is specific to pain (as opposed to being a marker of general sensory sensitivity).

The specificity of the model to pain was further characterized in light of the frequently reported weak association between self-perception of sensitivity to (intangible) pain and the actual pain thresholds during experimental stimuli (as measured via QST) (Edwards and Fillingim, 2007; Grundström et al., 2019; Meiselles et al., 2017; Ruscheweyh et al., 2009). In fact, self-reported measures have been shown to display a comparatively stronger associations with psychological measures (Coronado and George, 2018; Edwards and Fillingim, 2007), like anxiety or negative affect, rather than pain thresholds. Therefore, we investigated whether the proposed model is biased towards such psychological factors (anxiety, depression) and characterized the association of the predictions with various self-rated measures of pain. Additionally, to address this question with an increased statistical power, we tested potential model bias towards self-reported pain interference and pain intensity scores on cortical thickness data of (N=1095) participants from the Human Connectome Project. None of the analyses found any significant partial association with psychometrics or self-rated pain-related scores, which – especially together with the high power of the HCP dataset – strongly suggests that the predictions of proposed model do not primarily reflect one’s subjective self-evaluation of his or her own pain sensitivity but rather the actual pain thresholds, as assessed with behavioural responses to experimental pain.

The proposed predictive model’s biomedically relevant predictive performance, potential generalizability and high neuroscientific validity and specificity highlights that, next to functional brain signatures of pain (Spisak et al., 2020; Wager et al., 2013), brain morphology should also be considered a as promising modality for the objective brain-based characterisation of pain. The reported predictive performance, together the relatively short (approx. 3-4 minutes), reliable (Schwarz et al., 2016) and widely applicable data acquisition protocol and the lightweight data analysis requirements render brain morphology not only as an additional modality of future multimodal pain biomarkers (Tracey, 2021) - complementary to functional brain signatures of pain (Spisak et al., 2020; Wager et al., 2013) - but also as a highly accessible, standalone tool for pain research.

Next to serving as a proof-of-concept for the utility of morphology in the construction of pain-based predictive models, the present work also broadens our knowledge about the structural brain correlates of individual pain sensitivity differences. In all three regions that were robustly identified as valuable predictors (rACC, PHG and TP), we found that thicker cortex was associated with lower pain sensitivity. Cortical thickness has been shown to be systematically related to both cytoarchitecture and the structural hierarchical organisation of the cortex. Specifically, cortical thickness measures have been found to be inversely correlated with laminar differentiation (Wagstyl et al., 2015), myelination (Natu et al., 2019) and – in a region dependent manner – with neuronal density (la Fougère et al., 2011), altogether suggesting a close coupling between cortical structure and functional demand. To this end, while the mechanistic link between cortical thickness and the functional correlates of individual differences and alterations of pain sensitivity remains unclear, it is appealing to hypothesise that that the observed predictive capacity of cortical thickness in these regions originates from their differential functional involvement in pain processing, as a function of sensitivity to pain. Consistently with this notion, various studies have shown a significant association between cortical thickness and pain in healthy participants (Erpelding et al., 2012) as well as in pain disorders (DaSilva et al., 2007; Ellerbrock et al., 2013; Maleki et al., 2012)). The predictive regions identified in this study also fit well into this framework; the pain-related function of all these regions is widely acknowledged.

Thickness of the rACC was found to be the strongest predictor of pain sensitivity. In fact, the association of rACC thickness with pain sensitivity was significant even after correcting for multiple comparisons across all regions (via Bonferroni correction), suggesting that it may yield reasonable prediction performance on its own (r=-0.38). This finding fits well to the collective evidence for the involvement of rACC in modulating nociceptive processes (Apkarian et al., 2005; Bingel et al., 2008; Bingel et al., 2006; Eippert et al., 2009; Fuchs et al., 2014; Lamm et al., 2011; Smith et al., 2021; Spisák et al., 2017; Talbot et al., 1991; Wager et al., 2013). One could speculate that a thicker rACC could be a consequence of a higher functional involvement in nociception-related neural processes that are involved in attenuating pain. For example, Bingel et al. showed that repetitive pain stimulation over a number of days not only significantly decreased individual pain ratings, but also increased the functional involvement of rACC (Bingel et al., 2007). Furthermore, similarly to our finding, thicker rACC cortex has been linked to lower pain sensitivity levels in long-term meditation practitioners when compared to the control group (Grant et al., 2010). The lack of association between rACC thickness and self-reported pain intensity-interference in the analysis of large-scale cortical thickness data from HCP has interesting implications on rACC function in pain, especially considering the previously reported robust (N=505) divergence between QST-based pain thresholds and self-reported pain sensitivity (Edwards and Fillingim, 2007), with the latter being apparently more closely related to anxiety. This may suggest that the observed pain-related thickness changes in the rACC are specific to behaviour in the presence of actual pain experience (as measured with QST) and do not generalize to the subjective beliefs about one’s own sensitivity to (intangible) pain, as measured by self-reports.

The PHG and TP were also found to be robust predictors of pain sensitivity, although with a lower predictive coefficient and weaker unimodal association with pain sensitivity, as compared to rACC (r=-0.24 and -0.16, respectively). Several studies have described the involvement of PHG and hippocampal networks in nociceptive processes, (Bingel et al., 2002; Veldhuijzen et al., 2009) and its functional connectivity has been found to be an important modulator of pain quality experiences, possibly mediated by anticipatory anxiety and associative learning (Pinto et al., 2022). Several structural findings underpin the notion that differences in PHG function may manifest in regional morphology. For instance, Mutso et al. reported a reduction of hippocampal volume in chronic pain patients and proposed synaptic plasticity and neurogenesis as possible mechanisms, based on a neuropathic rodent pain model (Mutso et al., 2012). Moreover, in their aforementioned study, Grant et al. reported a negative association between cortical thickness and pain sensitivity in long-term mediation practitioners not only in the rACC, but interestingly, also in the PHG (Grant et al., 2010). Neumann et al. recently also reported increased grey matter volume in PHG with lower pain sensitivity (Neumann et al., 2021) in center 1 of the present multi-center analysis.

The predictive capacity of the thickness of the TP may also be traced back to its pain-related function. Several studies reported TP activation following noxious stimuli (Atlas et al., 2014; Godinho et al., 2006; Moulton et al., 2011; Shinozaki et al., 2016) and the TP has been brought into relation with the interaction of pain and working memory, changing the way stimuli are later remembered by acting directly on memory encoding (Godinho et al., 2006). The putative role of both the hippocampal formation and the TP in the formation of pain-related memories is underpinned by our results showing that morphological measures of these regions – but not the rACC - showed a weak but significant negative correlation with self-reported pain intensity and inference ratings in the HCP dataset. As self-reported pain scores are known to be associated to both anxiety (Edwards and Fillingim, 2007) and the interaction between pain and working memory (changing the way how painful experience is remembered) (Godinho et al., 2006), these results support the previously reported role of hippocampal and temporal formations in the exacerbation of pain by anxiety (Ploghaus et al., 2001).

## Conclusion

We have identified a structural MRI-based predictive model for estimating pain sensitivity across healthy individuals from multiple imaging centers. The robustness of the model across various cross-validation schemes, specificity to pain sensitivity and the neurological plausibility of the identified predictors demonstrate the capacity of brain morphology to predict individual sensitivity to pain. Our model is easy to apply on new data and might be a promising component of future multimodal imaging biomarker candidates of pain, with important implications for translational pain research and novel analgesic treatment strategies in precision medicine.

## Acknowledgement

This research was supported by the Deutsche Forschungsgemeinschaft (DFG, German Research Foundation): TRR 289 Treatment Expectation – project number 422744262.

