## Supplementary material for "Gray matter cortical thickness predicts individual pain sensitivity: a multi-center machine learning approach"

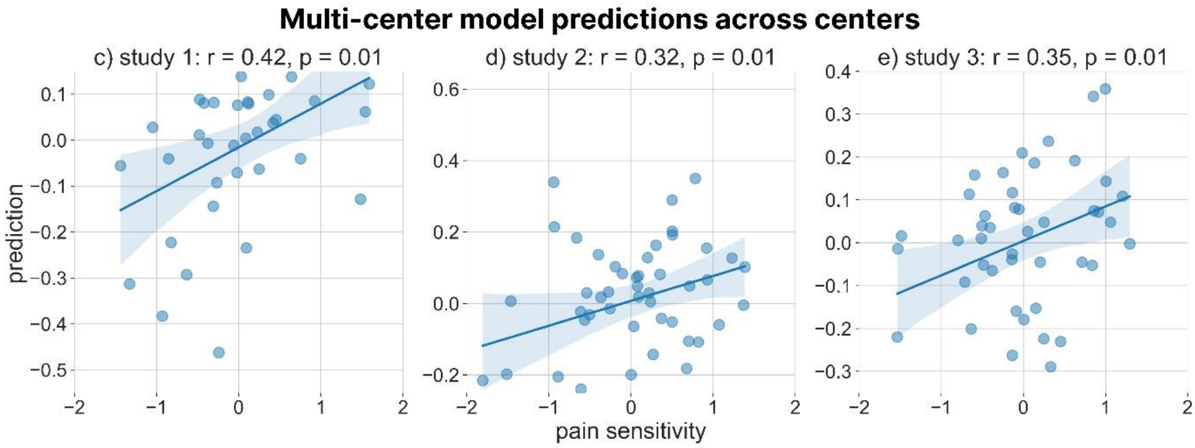


**Multi-center model predictions across centers**
**Supplementary figure 1**

Multi-center model predictions across three study centers are shown here (scatter plot with regression line: 95% confidence interval).

Supplementary table 1. Study specific additional variable details.

| No. | Additional measures | Study 1 (of N = 37) | Study 2 (N = 49) | Study 3 (N = 45) | Study 1 + 2 + 3) (N = 131) |
|---|---|---|---|---|---|
| **1** | BP MRI systole | - | 46 | 45 | 91 |
| **2** | BP MRI diastole | - | 46 | 45 | 91 |
| **3** | BP QST systole | - | 49 | 36 | 85 |
| **4** | BP QST diastole | - | 49 | 36 | 85 |
| **5** | MRI QST diff. | - | 49 | 45 | 94 |
| **6** | sex | 37 | 49 | 45 | 131 |
| **7** | day of menses | 10 | 23 | 25 | 58 |
| **8** | age | 37 | 49 | 45 | 131 |
| **9** | BMI | 37 | 49 | 45 | 131 |
| **10** | education | 37 | 48 | - | 85 |
| **11** | alcohol per unit | 37 | - | - | 37 |
| **12** | alcohol per week | 37 | 48 | - | 85 |
| **13** | PCS: catastrophizing | 34 | 49 | 29 | 112 |
| **14** | PCS: rumination | 37 | - | - | 37 |
| **15** | T50 | 37 | - | - | 37 |
| **16** | GLX | 37 | - | - | 37 |
| **17** | GABA | 37 | - | - | 37 |
| **18** | anxiety state | 36 | 49 | 28 | 113 |
| **19** | anxiety trait | 36 | 49 | 29 | 114 |
| **20** | PSQ | 36 | 49 | 29 | 114 |
| **21** | ADS-K | 36 | 49 | 28 | 113 |
| **22** | PSQ20 | - | 49 | 29 | 78 |
| **23** | PSQI | - | 49 | 23 | 72 |
| **24** | CDT | 37 | 49 | 45 | 86 |
| **25** | WDT | 37 | 49 | 45 | 86 |
| **26** | MDT | - | 49 | 45 | 49 |
| **27** | CPT | 37 | 49 | 45 | 131 |
| **28** | HPT | 37 | 49 | 45 | 131 |
| **29** | MPT (log transform) | 37 | 49 | 45 | 131 |
| **30** | batch | 37 | 49 | 45 | 131 |

Study specific list of additional measures are presented here, along with the number of participants for which these variables were available. '-' indicates that the additional measure is not present for that study.

Supplementary table 2. HCP1200 dataset: correlations with self-reported pain questionnaires

| region | Pain intensity (r) | p | Pain interference (r) | p |
|---|---|---|---|---|
| **Thickness** | | | | |
| **Left PHG** | 0.01 | 0.66 | 0.00 | 0.52 |
| **Right PHG** | 0 | 0.51 | 0.02 | 0.72 |
| **Left rACC** | 0 | 0.58 | -0.02 | 0.21 |
| **Right rACC** | 0.04 | 0.9 | 0.02 | 0.79 |
| **Left TP** | **-0.06** | **0.02** | -0.01 | 0.33 |
| **Right TP** | **-0.06** | **0.02** | -0.03 | 0.2 |
| **Area** | | | | |
| **Left PHG** | -0.04 | 0.1 | **-0.06** | **0.02** |
| **Right PHG** | -0.04 | 0.1 | **-0.06** | **0.02** |
| **Left rACC** | 0.01 | 0.59 | 0.01 | 0.6 |
| **Right rACC** | 0.02 | 0.79 | 0.01 | 0.64 |
| **Left TP** | -0.04 | 0.07 | -0.02 | 0.22 |
| **Right TP** | -0.03 | 0.17 | -0.03 | 0.15 |

Pearson’s r and its p-value are presented. P-value is generated using a non-parametric permutation testing with 10, 000 permutations.
